# IFT20 governs mesenchymal stem cell fate through positively regulating TGF-β-Smad2/3-Glut1 signaling mediated glucose metabolism

**DOI:** 10.1101/2022.02.28.482266

**Authors:** Yang Li, Shuting Yang, Yang Liu, Ling Qin, Shuying Yang

## Abstract

Aberrant lineage allocation of mesenchymal stem cells (MSCs) could cause bone marrow osteoblast-adipocyte imbalance, and glucose as an important nutrient is required for the maintenance of the MSCs’ fate and function. Intraflagellar transport 20 (IFT20) is one of the IFT complex B protein which regulates osteoblast differentiation, and bone formation, but how IFT20 regulates MSCs’ fate remains undefined. Here, we demonstrated that IFT20 controls MSC lineage allocation through regulating glucose metabolism during skeletal development. IFT20 deficiency in the early stage of MSCs caused significantly shortened limbs, decreased bone mass and significant increase in marrow fat. However, deletion of IFT20 in the later stage of MSCs and osteocytes just slightly decreased bone mass and bone growth and increased marrow fat. Additionally, we found that loss of IFT20 in MSCs promotes adipocyte formation, which enhances RANKL expression and bone resorption. Conversely, ablation of IFT20 in adipocytes reversed these phenotypes. Mechanistically, loss of IFT20 in MSCs significantly decreased glucose tolerance and suppressed glucose uptake and lactate and ATP production. Moreover, loss of IFT20 significantly inhibited TGF-β-Smad2/3 signaling and decreased the binding activity of Smad2/3 to *Glut1* promoter to downregulate Glut1 expression. These findings indicate that IFT20 plays essential roles for preventing MSC lineage allocation into adipocytes through controlling TGF-β-Smad2/3-Glut1 mediated glucose metabolism.

## Introduction

Bone marrow mesenchymal stem cells (MSCs) are multipotent stromal cells that display a transdifferentiation potential, are able to differentiate into cell lineages such as chondrocytes, osteoblasts, adipocytes, and myoblasts (Chen et al. 2016; Li and Wu 2020), and facilitate postnatal organ growth, repair and regeneration (Charbord 2010; Kalinina et al. 2011; Chen et al. 2016). As common progenitors, MSCs mobilize out of the bone marrow, migrate, and invade other tissues, such as bone or fat, to maintain bone homeostasis (Deng et al. 2011; Kumar and Ponnazhagan 2012). Accumulating evidence has shown a reciprocal interplay between adipogenesis and osteogenesis of MSCs in the bone marrow (Xiao et al. 2018; Yu et al. 2018; Deng et al. 2021). For instance, the factors that affect the balance between osteogenesis and adipogenesis within bone marrow could result in body composition changes (Schilling et al. 2014; Titorencu et al. 2014; Chen et al. 2016). Increased bone marrow adipose tissue (MAT) was observed in the context of aging and bone loss conditions (Justesen et al. 2001; Moerman et al. 2004). In addition, red bone marrow was reported to be gradually replaced by adipocytes during skeletal development (Jilka et al. 1999; Fan et al. 2017), and MAT accumulation increased at the expense of bone formation by inhibiting hematopoiesis and osteogenic regeneration (Ambrosi et al. 2017; Yu et al. 2018). Thus, disruption of the balance between adipogenesis and osteogenesis of MSCs may be a major cause of osteopenia and osteoporosis accompanied by progressive marrow adiposity. Nevertheless, the signaling molecules that control the temporal sequence of lineage allocation in MSCs are largely unknown.

MSCs are the progenitors of osteoblasts and adipocytes (James 2013; Yu et al. 2018). The adipogenesis and osteogenesis mediated by MSCs are the complicated processes, including the proliferation of MSC precursors, cell commitment to a specific lineage, and terminal differentiation (James 2013; Yu et al. 2018; Deng et al. 2021). In mammalian MSCs, peroxisome proliferator-activated receptor-gamma (PPARγ), CCAAT/enhancer-binding protein α (C/EBPα), fatty acid binding protein (Fabp4), and adiponectin are considered the key regulators of adipogenesis (Ghali et al. 2015; Ambele et al. 2016; Moseti et al. 2016), whereas alkaline phosphatase (ALP), runt-related transcription factor 2 (Runx2), osterix (OSX), and osteocalcin (OCN) are the main determinants of osteogenesis (Xiao et al. 2018; Li et al. 2021b). Although these transcription factors have been demonstrated to play essential roles in shifting MSC differentiation between adipocytes and osteoblasts, the underlying mechanisms that govern this differentiation commitment of MSCs remain undefined.

Studies have shown that glucose metabolism is a key hallmark of skeletal development (Karner and Long 2018; Lee et al. 2018). Glucose is a major source of energy in mammalian cells by generating ATP through intermediate glycolytic metabolites (Karner and Long 2018; Lee et al. 2018; Lee and Long 2018). In MSCs, glucose uptake was dramatically increased by upregulation of the expression of glycolytic enzymes such as hexokinase 2 (HK2), 6-phosphofructo-2-kinase/fructose-2,6-biphosphatase 3/4 (Pfkfb3/4), and lactate dehydrogenase A (LDHA) and the expression of glucose transporters (Gluts, particularly Glut1) to fuel aerobic glycolysis and provide cellular metabolites for the generation of new biomass, which ultimately stimulates bone formation (Dirckx et al. 2018; Lee and Long 2018). It’s well-known that TGF-β signaling is a critical regulator of glucose metabolism during bone development and homeostasis (Kitagawa et al. 1991; Andrianifahanana et al. 2016; Wu et al. 2016; Xu et al. 2018). However, how TGF-β signaling is regulated for controlling glucose metabolism during skeletal development and marrow osteoblast and adipocyte homeostasis is largely unknown.

IFT proteins play critical roles in the bidirectional transport of molecules along cilia and are essential for cilia formation and function (Yuan et al. 2016; Lim et al. 2020; Saternos et al. 2020). Disruption of primary cilia and ciliary protein results in abnormal skeletal development and remodeling (Yuan et al. 2016; Xiao et al. 2018; Lim et al. 2020). Mutations of cilia-related proteins, such as Bardet-Biedl syndrome (BBS), alström syndrome 1 (ALMS1), and IFT88, can cause obesity with abnormal glucose metabolism (Oh et al. 2015; Vaisse et al. 2017; Lee et al. 2020). Our previous study showed IFT80 promoted chondrogenesis and fracture healing through regulation of TGF-β-Smad2/3 signaling (Liu et al. 2020). Other studies have showed that loss of IFT88 protein in MSCs could inhibit TGF-β signaling (Labour et al. 2016) and canonical TGF-β-Smad2/3 activation can upregulate Glut1 expression (Andrianifahanana et al. 2016). However, it remains unknown whether IFT proteins are critical regulator of MSC lineage commitment through controlling glucose metabolism during bone development. IFT20 is one of the IFT complex B proteins that expresses in many tissues and cells including MSCs, and regulates cilia and bone formation (Lim et al. 2020; Yang et al. 2021), but how IFT20 regulates MSCs’ fate is undefined. In this study, we characterized the contribution of IFT20 to the proliferation, differentiation, and maturation of MSCs and determined the molecular mechanisms of IFT20-driven MSCs’ fate. Our data revealed that IFT20 controls MSC lineage allocation by regulating TGF-β-Smad2/3-Glut1-mediated glucose metabolism.

## Results

### Loss of IFT20 in MSCs causes significantly shortened limbs and inhibits skeletal development

Prx1-Cre is expressed throughout the limb bud stage from embryonic Day 9.5 (E9.5) (Yu et al. 2019) and is expressed in chondrocytes and during all stages of osteoblast differentiation (Yu et al. 2018; Yu et al. 2019; Deng et al. 2021). To investigate the function of IFT20 in MSCs, we first generated an IFT20 conditional knockout mouse model by crossing IFT20^f/f^ mice with Prx1-Cre mice (hereafter named Prx1-Cre;IFT20^f/f^ mice) in which IFT20 was deleted in the early stage of MSCs. qRT-PCR data verified that IFT20 was largely abrogated in bone marrow-derived MSCs (Supplemental Fig. S1A). We found that the vertebrate bone length was slightly shorter but that the limbs were significantly shorter in the Prx1-Cre;IFT20^f/f^ mice than those in the control littermates (Fig. 1A). Newborn Prx1-Cre;IFT20^f/f^ mice displayed impaired intramembranous ossification with hypomineralization of the calvarium (Fig. 1B). Notably, the limbs were devoid of bone and the rib cage was disturbed in the Prx1-Cre;IFT20^f/f^ newborns compared to the age-matched controls (Fig. 1C, D). To further investigate the potential contribution of IFT20 to skeletal development, we next evaluated the skulls, tibiae and femurs at different developmental stages of the Prx1-Cre;IFT20^f/f^ embryos (E16.5 and E18.5). Whole-mount skeletal staining showed that all limbs and skull bone were notably undermineralized. Additionally, the limbs were significantly shorter in embryonic and 1-month-old Prx1-Cre;IFT20^f/f^ mice compared with age-matched controls (Supplemental Fig. S1B-F), suggesting IFT20 in the limb mesenchyme is necessary for skeletal development in the prenatal and newborn stages.

**Figure 1.**
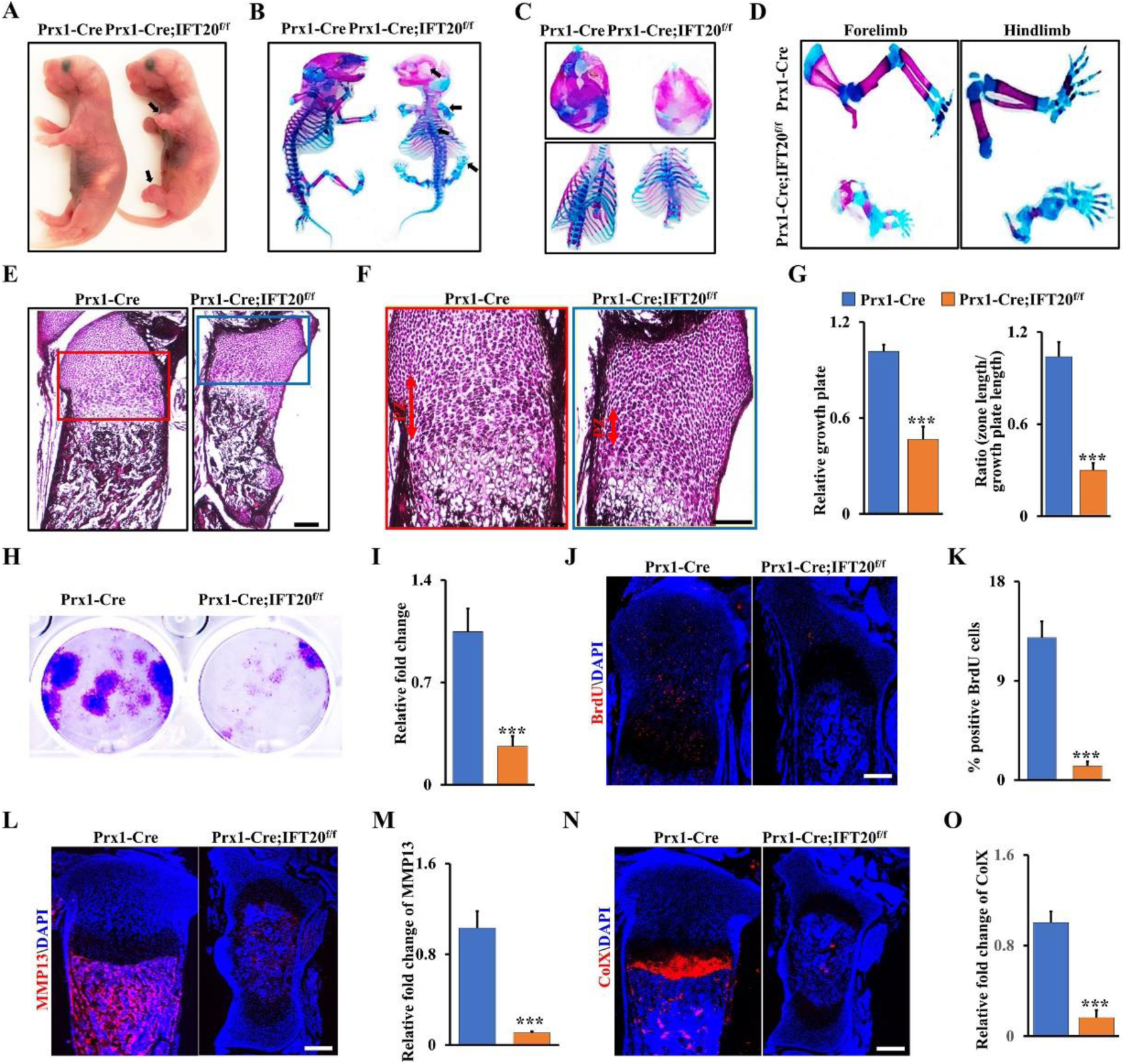
Loss of IFT20 in MSCs causes significantly shortened limbs and inhibits skeletal development. (A-D) Representative whole-mount skeletal-stained image of Prx1-Cre;IFT20^f/f^ mice and age-matched controls at the newborn stage (P0). The black arrows direct to the shorten limbs and the serious area of bone loss at P0. (E) Representative H&E-stained image of the tibiae from Prx1-Cre;IFT20^f/f^ mice and controls at P0. Scale bars, 100 μm. (F) High magnification image of box area from (E). Scale bars, 50 μm. (G) Relative length of growth plate and the proliferation zone (PZ) was identified based on (E, F) as indicated. (H, I) Representative image of colony formation (CFU) stained with 0.5% crystal violet, and quantified colony numbers as indicated. (J, K) Representative fluorescence image of BrdU^+^ in the tibiae from Prx1-Cre;IFT20^f/f^ mice and controls at P0. Scale bars, 100 μm. The BrdU^+^ cells were quantified in the corresponding column (K). (L-O) Representative fluorescence image of MMP13 and ColX at P0 tibiae, and the corresponding quantification as indicated. Error bars were the means ± SEM from three independent experiments. \*\*\**P* < 0.001.

Given that IFT20 deficiency in Prx1-expressing cells impaired bone formation with severe shorten limb in mice, we further examined whether IFT20 deficiency could inhibit chondrocyte formation and maturation. Close histologic examination of the tibiae revealed that the proliferation zone (PZ) in the growth plate of the Prx1-Cre;IFT20^f/f^ mice was shortened, and there was an apparent defect in chondrocyte hypertrophy compared to that in the age-matched controls (Fig. 1E-G). Consistently, colony-forming unit (CFU) assays confirmed that the proliferation rate of MSCs was greatly decreased after deletion of IFT20 (Fig. 1H, I). Concomitantly, BrdU labeling analyses showed few proliferating cells in the growth plate of tibiae from the Prx1-Cre;IFT20^f/f^ newborns compared to the age-matched controls (Fig. 1J, K). Moreover, the expression levels of the hypertrophic chondrocyte marker collagen X (ColX) and the terminal hypertrophy marker matrix metallopeptidase 13 (MMP13) were dramatically decreased in chondrocytes from the Prx1-Cre;IFT20^f/f^ newborns (Fig. 1L-O), suggesting that deletion of IFT20 with Prx1-Cre impairs chondrocyte proliferation and maturation. Overall, our data showed IFT20 in MSCs plays a critical role during skeletal development.

### IFT20 deficiency in MSCs causes bone loss and MAT accumulation

Micro-CT analysis of femurs and skull bones showed a marked decrease in mineralized tissues (Fig. 2A, B and Supplemental Fig. S2). The femurs from the Prx1-Cre;IFT20^f/f^ mice lost approximately 50% of bone volume per total volume (BV/TV), 20% of trabecular thickness (Tb.Th), and 30% of the trabecular number (Tb.N), and trabecular separation (Tb.Sp) showed a 1.51-fold increase compared to those in the controls (Fig. 2C). Consistently, the trabecular volumetric bone mineral density (BMD) and serum OCN level also dramatically decreased compared to that in the age-matched group (Fig. 2D, E). Interestingly, accompanied by a reduction in bone mass, we found an apparent increase in MAT accumulation and a 6.6-fold increase in adipocyte numbers in the Prx1-Cre;IFT20^f/f^ mice compared to the age-matched controls (Fig. 2F-H), as evidenced by osmium tetroxide (OsO_4_) staining and micro-CT analysis (Fig. 2I), thereby supporting the histological findings (Fig. 2F, G). Additionally, histomorphometric analysis of the femur metaphysis demonstrated that deletion of IFT20 decreased the dynamic indices of the bone formation rate (BFR) and bone formation mineral apposition rate (MAR) in the Prx1-Cre;IFT20^f/f^ mice (Fig. 2J-L). Notably, we also observed higher osteoclast activity in the Prx1-Cre;IFT20^f/f^ mice by tartrate-resistant acid phosphatase (TRAP) staining of sections from the femur (Fig. 2M).

**Figure 2.**
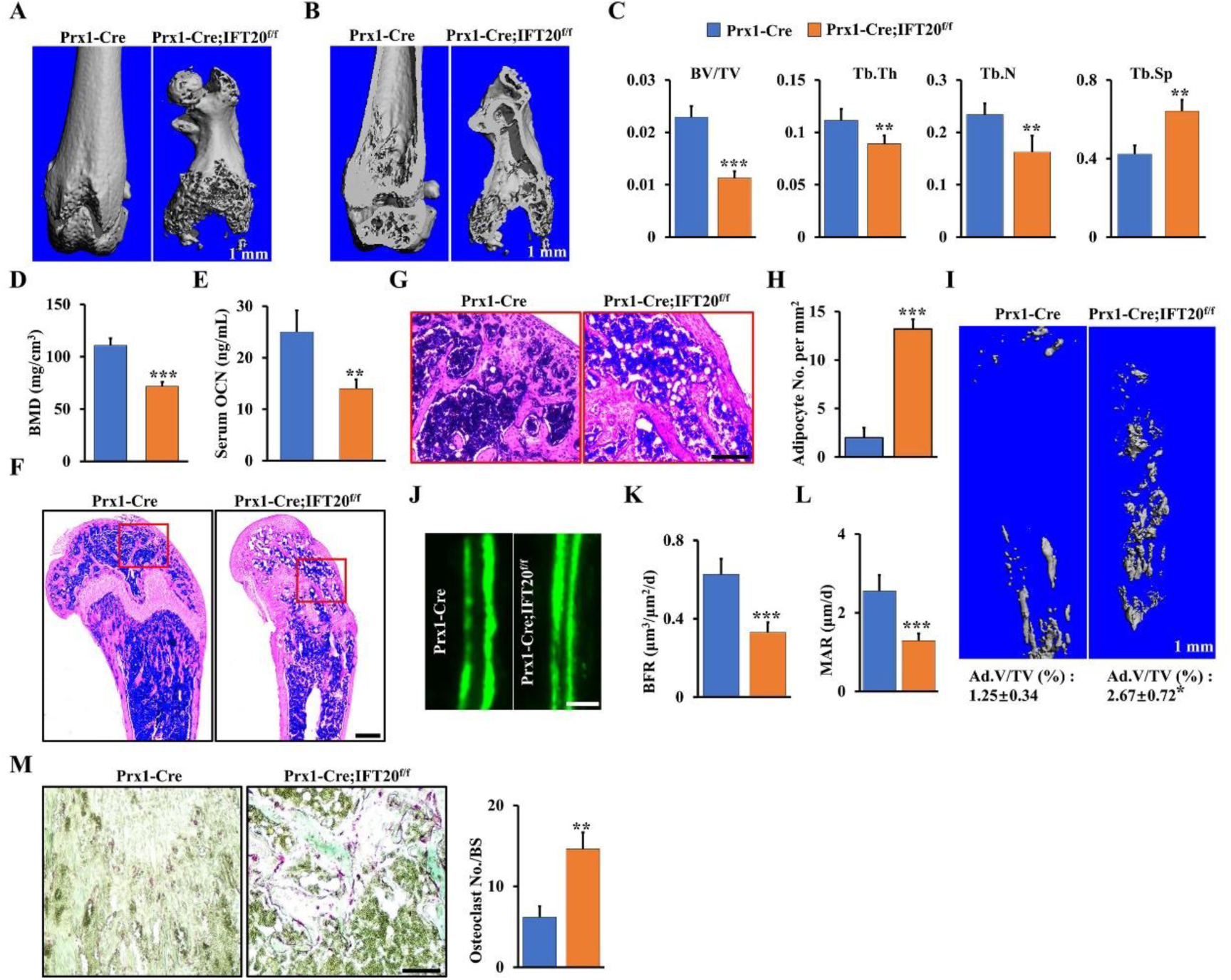
IFT20 deficiency in MSCs causes bone loss and MAT accumulation. (A, B) Representative micro-CT image of femurs of Prx1-Cre;IFT20^f/f^ mice and controls at 1 month. Scale bars, 1 mm. (C) Histomorphometric analysis of bone parameters in the femurs of 1-month-old Prx1-Cre;IFT20^f/f^ mice and controls. Bone volume fraction (BV/TV); trabecular thickness (Tb.Th); trabecular number (Tb.N); trabecular spacing (Tb.Sp). N=5 mice/group. (D) Quantitative measurements of bone mineral density (BMD) of femurs from Prx1-Cre;IFT20^f/f^ mice and controls at 1 month. (E) The serum level of OCN from Prx1-Cre;IFT20^f/f^ mice and controls at 1 month. (F) Representative H&E-stained image of femur sections from 1-month-old Prx1-Cre;IFT20^f/f^ mice and controls. Scale bars, 200 μm. (G) High magnification image of red box area from (F). Scale bars, 100 μm. (H) Adipocyte numbers per tissue area were identified based on the H&E images of (F). (I) OsO_4_ micro-CT staining of decalcified tibiae by micro-CT analysis as indicated. (J-L) Calcein double labeling in tibia of 1-month-old Prx1-Cre;IFT20^f/f^ mice and controls. Scale bar, 50 μm. (M) Representative TRAP-stained image of femur sections from 1-month-old Prx1-Cre;IFT20^f/f^ mice and controls. Scale bar, 100 μm. The corresponding quantitative analysis of TRAP staining were at right. Error bars were the means ± SEM from three independent experiments. \**P* < 0.05, \*\**P* < 0.01, \*\*\**P* < 0.001.

Recently, leptin receptor (Lepr) was reported to be a marker of MSCs at the late stage behind Prx1 and drives the differentiation of osteoblasts and adipocytes in adult mice (Kfoury and Scadden 2015; Yue et al. 2016; Yang et al. 2019). To further validate whether IFT20 deficiency at the late stage of MSCs causes bone loss and MAT accumulation, we next generated Lepr-Cre;IFT20^f/f^ mice. Consistently, the Lepr-Cre;IFT20^f/f^ mice recapitulated the osteogenic inhibition that was observed in the Prx1-Cre;IFT20^f/f^ mice (Supplemental Fig. S3). Micro-CT analysis of femurs showed a significant reduction in trabecular bone mass in the Lepr-Cre;IFT20^f/f^ mice compared to the age-matched controls (Supplemental Fig. S3A-G). Dynamic histomorphometric analysis revealed a significant inhibition of bone formation in the Lepr-Cre;IFT20^f/f^ mice (Supplemental Fig. S3H-J), suggesting that deletion of IFT20 in later stage of MSCs could lead to impaired bone formation in mice, as evidenced by H&E and TRAP staining (Supplemental Fig. S3K, L). Unexpectedly, even though bone formation was significantly blocked, there was no marked difference in adipocyte numbers between the Lepr-Cre;IFT20^f/f^ mice and the age-matched controls (Supplemental Fig. S3K). The results were further confirmed by OsO_4_ micro-CT analyses (Supplemental Fig. S3M), suggesting that IFT20 may direct lineage decisions at the early stage of MSCs. To further corroborate our hypothesis, we generated osteocyte-specific knockout mice (DMP1-Cre;IFT20^f/f^). As expected, we found that IFT20 was dispensable in osteocytes (Supplemental Fig. S4). Thus, these findings indicated that IFT20 governs the bone-fat balance by directing the lineage decisions of MSCs at an early stage.

### Adipocytes are the principal source for RANKL after loss of IFT20 in MSCs, and IFT20 deficiency in adipocytes reverses bone phenotype by reducing RANKL expression

Bone marrow adipocytes have been reported to secrete receptor activator of NF-κB ligand (RANKL) and support osteoclast differentiation (Boyce and Xing 2007; Park et al. 2017; Yu et al. 2021). Our data showed that deletion of IFT20 in the mesenchymal linage increased osteoclastogenesis. To further explore whether IFT20 affects the expression and function of RANKL in MSCs, we performed histological analysis of the long bone and marrow cavity. We found that many adipocytes were present in very close proximity to the bone surface and covered by TRAP^+^ osteoclasts in the Prx1-Cre;IFT20^f/f^ mouse femurs compared to the controls (Fig. 3A). To determine whether increased osteoclastogenesis results from the alteration of RANKL in the Prx1-Cre;IFT20^f/f^ mice, we isolated RNA from the whole bone marrow and bone marrow adipose tissue to detect RANKL expression. We found that RANKL expression was significantly upregulated in both whole bone marrow and bone marrow adipose tissue after deletion of IFT20 (Fig. 3B, C). Interestingly, the serum levels of RANKL from the Prx1-Cre;IFT20^f/f^ mice showed a marked elevation (Fig. 3D). Moreover, a remarkably increased ratio of RANKL/osteoprotegerin (OPG) was observed in the serum of the Prx1-Cre;IFT20^f/f^ mice compared to the control mice, suggesting that IFT20 may play a role in RANKL secretion (Fig. 3E).

**Figure 3.**
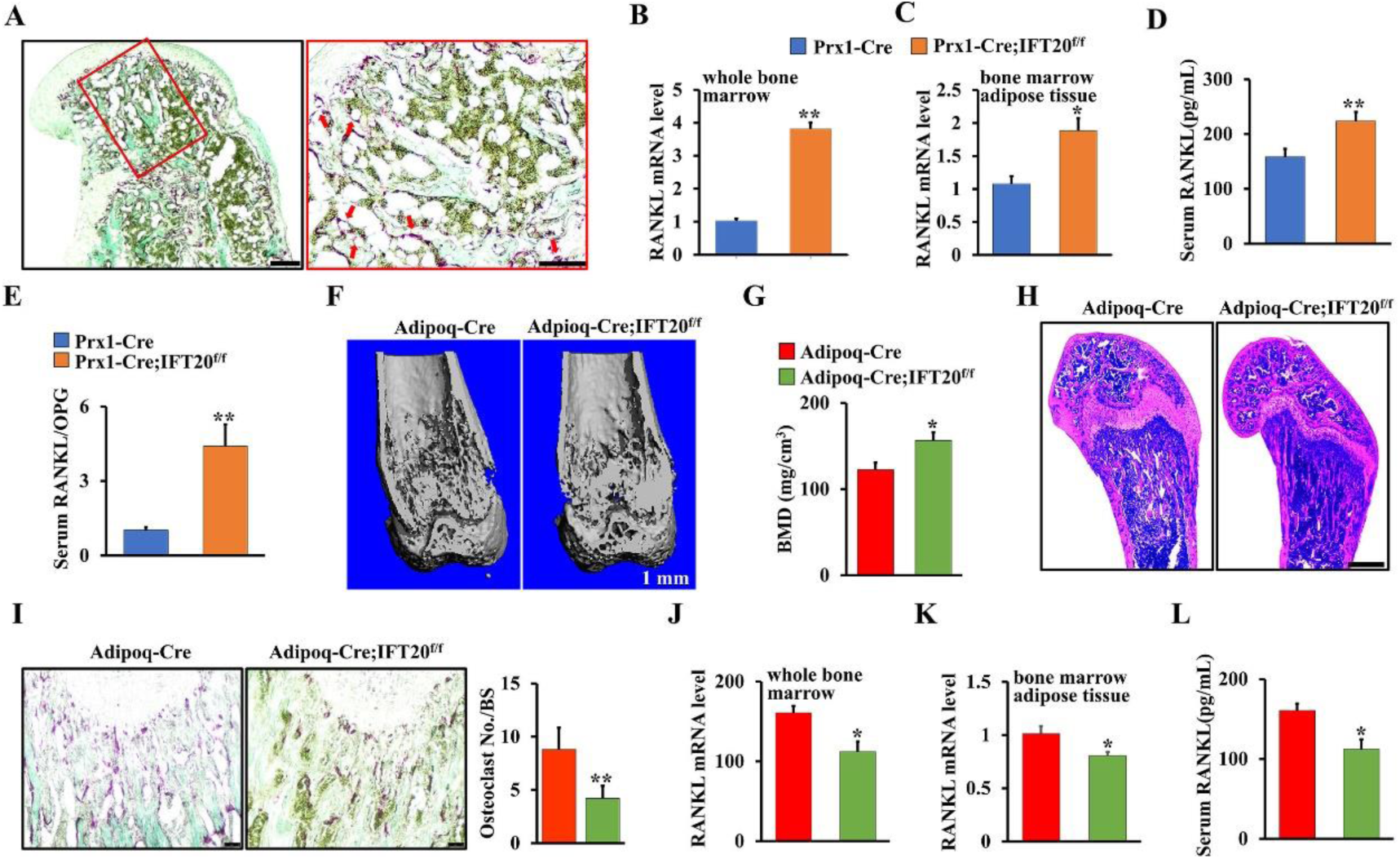
Adipocytes are the principal source for RANKL after loss of IFT20 in MSCs, and IFT20 deficiency in adipocytes reverses bone phenotype by reducing RANKL expression. (A) Representative TRAP-stained image of femurs from 1-month-old Prx1-Cre;IFT20^f/f^ mice and controls. Scale bar, 200 μm. High magnification image of red box area was at right. Scale bars, 100 μm. The red arrow indicates adipocytes that are found in close vicinity to TRAP positive osteoclasts. (B, C) qRT-PCR analysis of RANKL in whole bone marrow (B) and bone marrow adipose tissue (C). (D) The serum level of RANKL from 1-month-old Prx1-Cre;IFT20^f/f^ mice were significantly increased compared to age-mated controls. (E) The serum RANKL/OPG ratio was identified in 1-month-old Prx1-Cre;IFT20^f/f^ mice and controls as indicated. (F) Representative micro-CT image of femurs from 1-month-old Adipoq-Cre;IFT20^f/f^ mice and controls. Scale bars, 1 mm. (G) Quantitative BMD measurements of femurs from 1-month-old Adipoq-Cre;IFT20^f/f^ mice and controls. (H) Representative H&E-stained image of femurs from 1-month-old Adipoq-Cre;IFT20^f/f^ mice and controls. Scale bars, 200 μm. (I) Representative TRAP-stained image of femurs from 1-month-old Adipoq-Cre;IFT20^f/f^ mice and controls. Scale bar, 100 μm. The corresponding quantitative analysis of TRAP staining were at right. (J, K) qRT-PCR analysis of RANKL in whole bone marrow (J) and bone marrow adipose tissue (K). (L) The serum levels of RANKL from 1-month-old Adipoq-Cre;IFT20^f/f^ mice were significantly decreased compared to controls. Error bars were the means ± SEM from three independent experiments. \**P* < 0.05, \*\**P* < 0.01.

Since deletion of IFT20 in MSCs prohibited bone formation but promoted marrow adipogenesis, we explored whether deletion of IFT20 in adipocytes using adipocyte-specific knockout mice (Yu et al. 2021) (hereafter named Adipoq-Cre;IFT20^f/f^) could reverse the bone-fat imbalance that was observed in the Prx1-Cre;IFT20^f/f^ mice. As expected, micro-CT analysis showed that there was a significant increase in bone mass but no MAT accumulation in trabecular bones from the 1-month-old Adipoq-Cre;IFT20^f/f^ mice compared with the controls (Fig. 3F-H). TRAP staining analysis showed that osteoclast formation was significantly suppressed after deletion of IFT20 in adipocytes (Fig. 3I). qRT-PCR analysis of the whole bone marrow and bone marrow adipose tissue demonstrated that the deletion of IFT20 in adipocytes inhibited RANKL expression (Fig. 3J, K). To further confirm this, we also investigated the serum RANKL level in the Adipoq-Cre;IFT20^f/f^ mice and controls by ELISAs. Consistently, serum RANKL was significantly decreased in the Adiopq-Cre;IFT20^f/f^ mice compared to the age-matched controls (Fig. 3L), indicating that RANKL may be mainly derived from marrow adipocytes in Prx1-Cre;IFT20^f/f^ mice.

### Deletion of IFT20 in MSCs promotes adipocyte differentiation but inhibits osteoblast differentiation

To further characterize the function of IFT20 in MSCs’ fate, we isolated MSCs from the Prx1-Cre;IFT20^f/f^ and control mice respectively and identified the effect of IFT20 on osteogenesis and adipogenesis *in vitro*. Deletion of IFT20 in MSCs resulted in a significant decrease in alkaline phosphatase (ALP)-positive cells (Fig. 4A) and mineralized nodule formation (ARS) (Fig. 4B) after stimulation with osteogenic media for 5 and 14 days, respectively. Moreover, we found that deletion of IFT20 in MSCs decreased ALP activity (Fig. 4C), suggesting that IFT20 is essential for osteoblast differentiation and bone formation. In contrast, Oil Red O staining results showed a marked increase in adipogenesis from IFT20-deficient MSCs after stimulation with adipogenic media for 14 days (Fig. 4D). Hence, these results suggested that IFT20 governs the balance of osteogenesis and adipogenesis from MSCs. To further confirm the function of IFT20 in commitment of MSCs, we performed qRT-PCR after stimulation with osteogenic or adipogenic media for 14 days. The results showed that deletion of IFT20 in MSCs significantly downregulated the expression of osteogenic markers (ALP, Runx2, OSX and OCN) but upregulated the expression of adipogenic genes (PPARγ, C/EBPα, Fabp4, and adiponectin) compared to those in the control cells (Fig. 4E, F). These data demonstrated that IFT20 controls osteogenic and adipogenic differentiation of skeletal stem cells.

**Figure 4.**
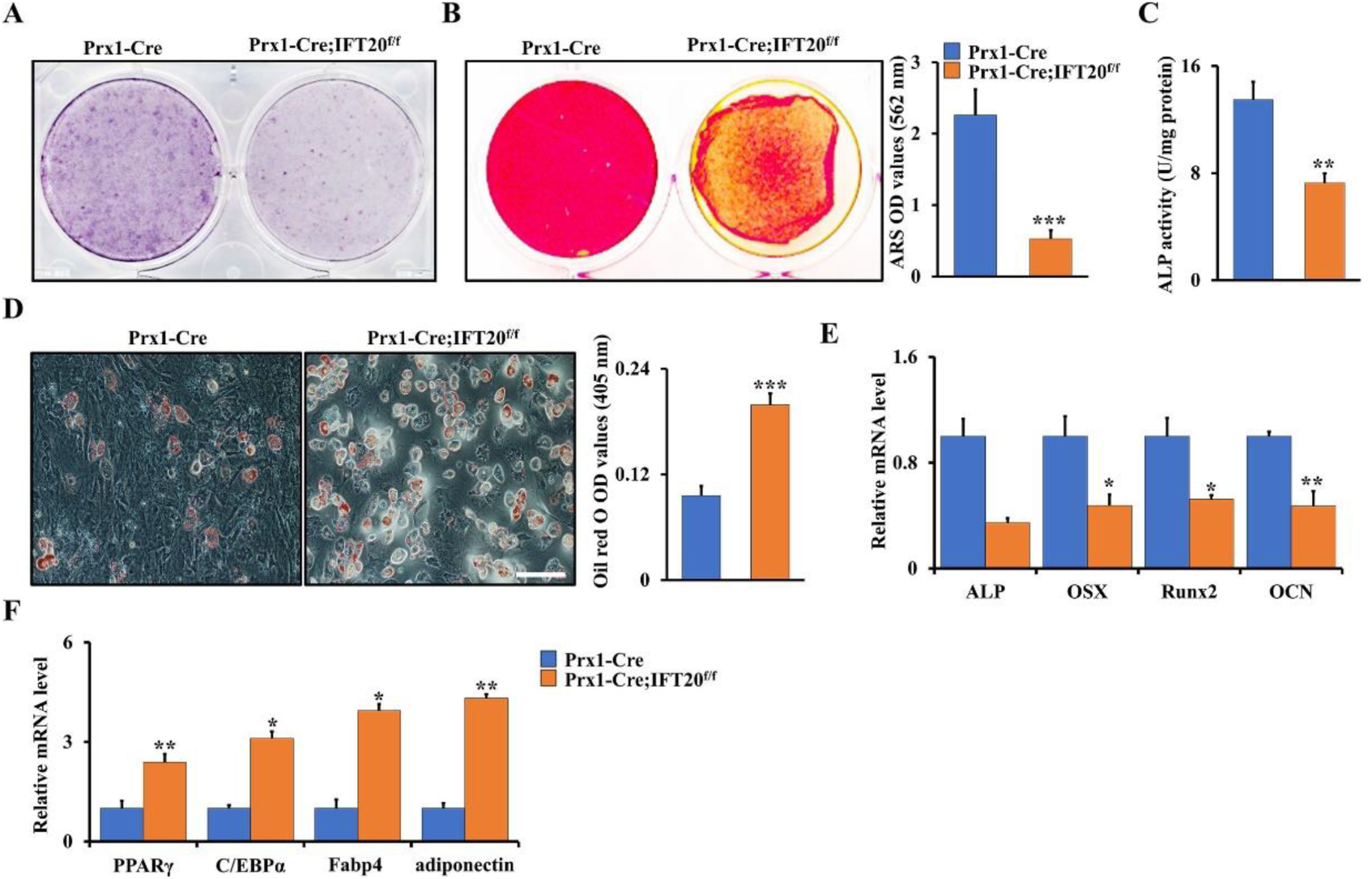
Deletion of IFT20 in MSCs promotes adiopogenic differentiation but inhibits osteogenic differentiation. (A) Representative image of ALP (Day 5) staining of MSCs from Prx1-Cre;IFT20^f/f^ and littermate control mice. (B) Representative image of ARS (Day 14) staining of MSCs from Prx1-Cre;IFT20^f/f^ and littermate control mice. Relative OD of ARS staining was at right. (C) ALP activity. (D) Representative image of Oil red O staining after adipogenic incubation of 14 days. The corresponding quantitative analyses of Oil red O staining were performed at right. (E, F) Osteogenic and adipogenic markers analyses by qRT-PCR after stimulation with osteogenic or adipogenic media for 14 days as indicated. Error bars were the means ± SEM from three independent experiments. \**P* < 0.05, \*\**P* < 0.01, \*\*\**P* < 0.001.

### Deletion of IFT20 in MSCs decreases glucose tolerance which is restored by deletion of IFT20 in adipocytes

Glucose is the main energy source in skeletal development, which is regulated by Glut family proteins such as Glut1-4, and Glut1 has been proven to be the most important glucose transporter in osteoblasts (Karner and Long 2018; Lee et al. 2018). Previous studies demonstrated that cilia-related proteins contribute to regulation of obesity and energy metabolism, and dysfunction of cilia causes metabolic defects (Oh et al. 2015; Vaisse et al. 2017; Lee et al. 2020). To further investigate whether deletion of IFT20 in MSCs affects glucose metabolism, we generated a tamoxifen-inducible IFT20 conditional knockout mouse model in which IFT20 was specifically deleted in MSCs (hereafter named Prx1-Cre^ERT^;IFT20^f/f^) at postnatal stage. By examining the blood glucose and insulin levels in 1-month-old Prx1-Cre^ERT^;IFT20^f/f^ mice and controls under random-fasted conditions, we found a significant increase in blood glucose in the Prx1-Cre^ERT^;IFT20^f/f^ mice compared to the controls; however, there was no significant change in serum insulin (Fig. 5A, B). Consistently, blood glucose levels were also increased in the Prx1-Cre;IFT20^f/f^ mice under random feeding conditions from P1 to P28 (Fig. 5C). Moreover, the levels of triglycerides and blood glucose in the 1-month-old Prx1-Cre;IFT20^f/f^ mice were increased under random-fasted conditions compared to those in the Cre control mice (Fig. 5D, E). To further explore IFT20 regulation of glucose, we carried out glucose tolerance tests following intraperitoneal injection of glucose in the Prx1-Cre;IFT20^f/f^ mice and the control mice. Our data showed impaired glucose tolerance in the Prx1-Cre;IFT20^f/f^ mice (Fig. 5F). Moreover, we found that the level of glycogen stored in the liver was increased due to IFT20 deficiency in MSCs (Fig. 5G, H). Insulin is a well-known anabolic hormone that maintains the uptake and content of glucose in the serum (Nguyen et al. 2011; Cipriani et al. 2020). Our results showed no pronounced alterations in serum insulin in the Prx1-Cre;IFT20^f/f^ mice under either fed or random-fasted conditions (Fig. 5I, J), as evidenced by an insulin secretion test after glucose injection (Fig. 5K), suggesting that IFT20 in MSCs inhibits glucose metabolism but does not affect insulin levels. Moreover, the defect in glucose metabolism appeared to be reversed after deletion of IFT20 in adipocytes using Adipoq-Cre (Fig. 5L, M), suggesting that IFT20 governs lineage allocation of mesenchymal stem cells by regulating glucose metabolism.

**Figure 5.**
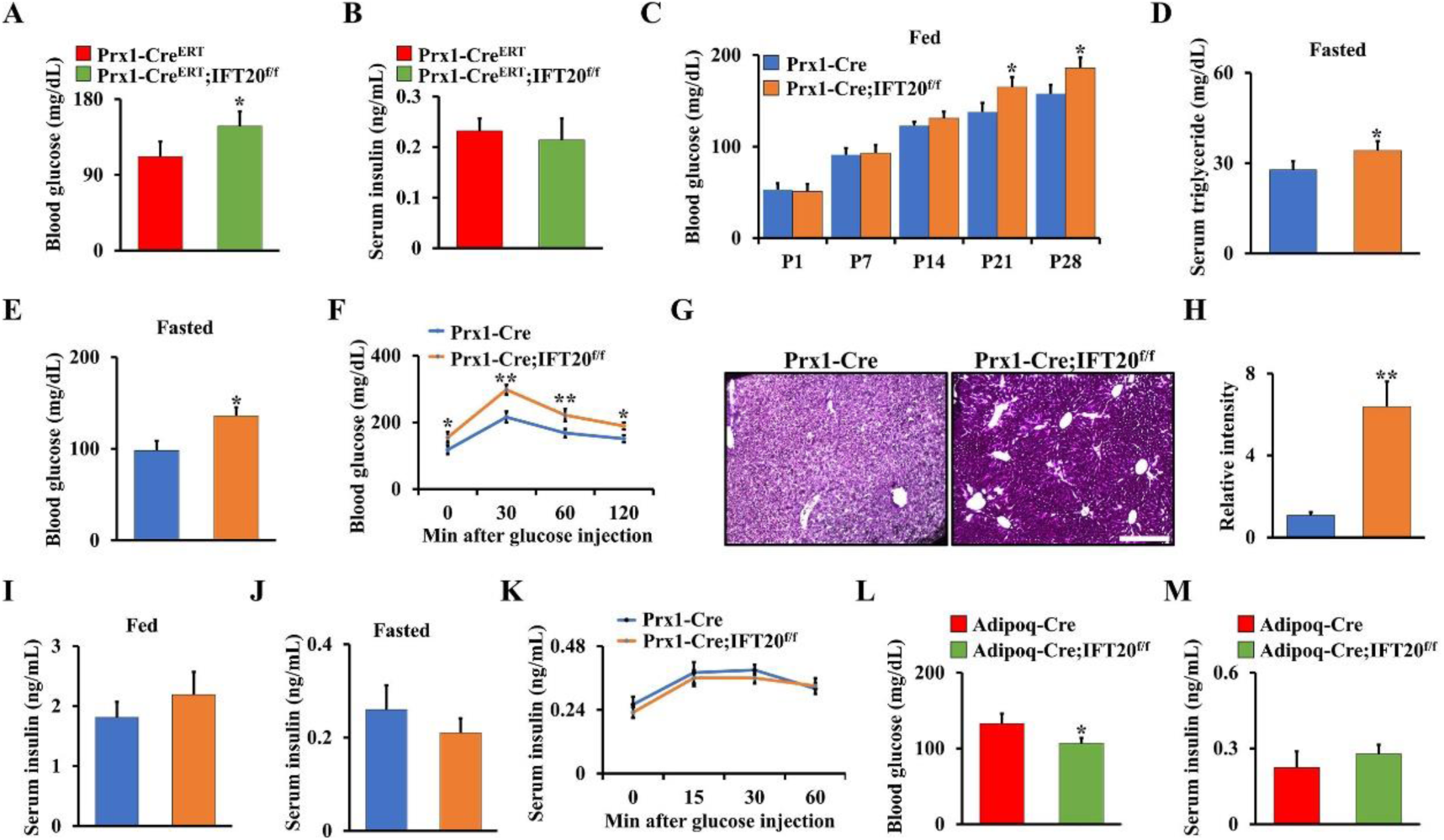
Deletion of IFT20 in MSCs decreases glucose tolerance which is restored by deletion of IFT20 in adipocytes. (A, B) Blood glucose (A) and insulin (B) levels after overnight fasting at 1-month-old Prx1-Cre^ERT^;IFT20^f/f^ mice and controls. N=6. (C) Blood glucose levels in random-fed state from P1 to P28. N=8 per group. (D, E) The levels of blood triglyceride (D) and glucose (E) after overnight fasting at 1-month-old mice as indicated. N=6. (F) Glucose tolerance test. (G) PAS staining on liver as indicated, revealing glycogen content. (H) The corresponding quantitative intensity of PAS staining. (I, J) Serum insulin levels in random-fed (I) and -fasted (J) conditions. (K) glucose-stimulated insulin secretion test. (L, M) Blood glucose (L) and insulin (M) levels after overnight fasting at 1-month-old Adipoq-Cre;IFT20^f/f^ mice and controls. N=6. Error bars were the means ± SEM from three independent experiments. \**P* < 0.05, \*\**P* < 0.01.

### IFT20 promotes glucose uptake and glycolysis in MSCs through Glut1 signaling

Previous findings showed Glut1-mediated glucose metabolism is essential for skeletal development (Karner and Long 2018; Lee et al. 2018; Wang et al. 2021). To explore whether IFT20 affects glucose metabolism through regulation of Glut1. We first examined the expression pattern of Glut1 during skeletal development. Our data showed that Glut1 expression significantly decreased in chondrocytes from the Prx1-Cre;IFT20^f/f^ embryos (E14.5, E16.5 and E18.5) compared to their age-matched controls (Fig. 6A). Additionally, Glut1 expression was inhibited in the 1-month-old Prx1-Cre;IFT20^f/f^ animals compared to the controls (Fig. 6B, C). To further explore glucose metabolism in the Prx1-Cre;IFT20^f/f^ mice, we evaluated the expression of key genes encoding glycolysis-regulating enzymes, such as Glut1-4, HK2, Pfkfb3/4, and Ldha. Our data suggested that these genes, especially Glut1, had significantly downregulated expression in MSCs from the Prx1-Cre;IFT20^f/f^ mice compared to the controls (Fig. 6D-F). To corroborate the above observations and further assess the role of IFT20 in regulating glucose metabolism, we isolated primary MSCs from IFT20^f/f^ mice and infected them with adenoviruses expressing Cre (Ad-Cre) or the control (Ad-GFP) *in vitro*. qRT-PCR analyses showed the deletion of IFT20 in the Ad-Cre-treated cells (Supplemental Fig. S5A). Consistently, the IFT20-deficient MSCs showed a decrease in the expression of Glut1-4, especially Glut1 (Fig. 6G). Additionally, the IFT20-deficient MSCs showed decreased glucose uptake and lactate and ATP production, as documented by analysis of conditioned culture media (Fig. 6H-J). Quantification of cellular uptake of 2-NBDG, a fluorescently labeled glucose analog, demonstrated a significant decrease in the IFT20-deficient MSCs compared to the control MSCs with Ad-GFP transduction (Fig. 6K), indicating that IFT20 governs Glut1-mediated glucose metabolism in MSCs. To visualize glucose uptake *in vivo*, we treated 1-month-old Prx1-Cre;IFT20^f/f^ mice and controls with the fluorescent glucose analog 2-NBDG and then assessed them 45 minutes after the injection. By evaluating the uptake and accumulation of the compound in the tibiae, we found that glucose uptake was significantly inhibited by loss of IFT20 in MSCs (Fig. 6L). More importantly, glucose metabolism could be reversed after forced expression of IFT20 in MSCs (Fig. 6M-O and Supplemental S5B). Overall, our data suggest that IFT20 directs MSC fate through Glut1-mediated glucose metabolism.

**Figure 6.**
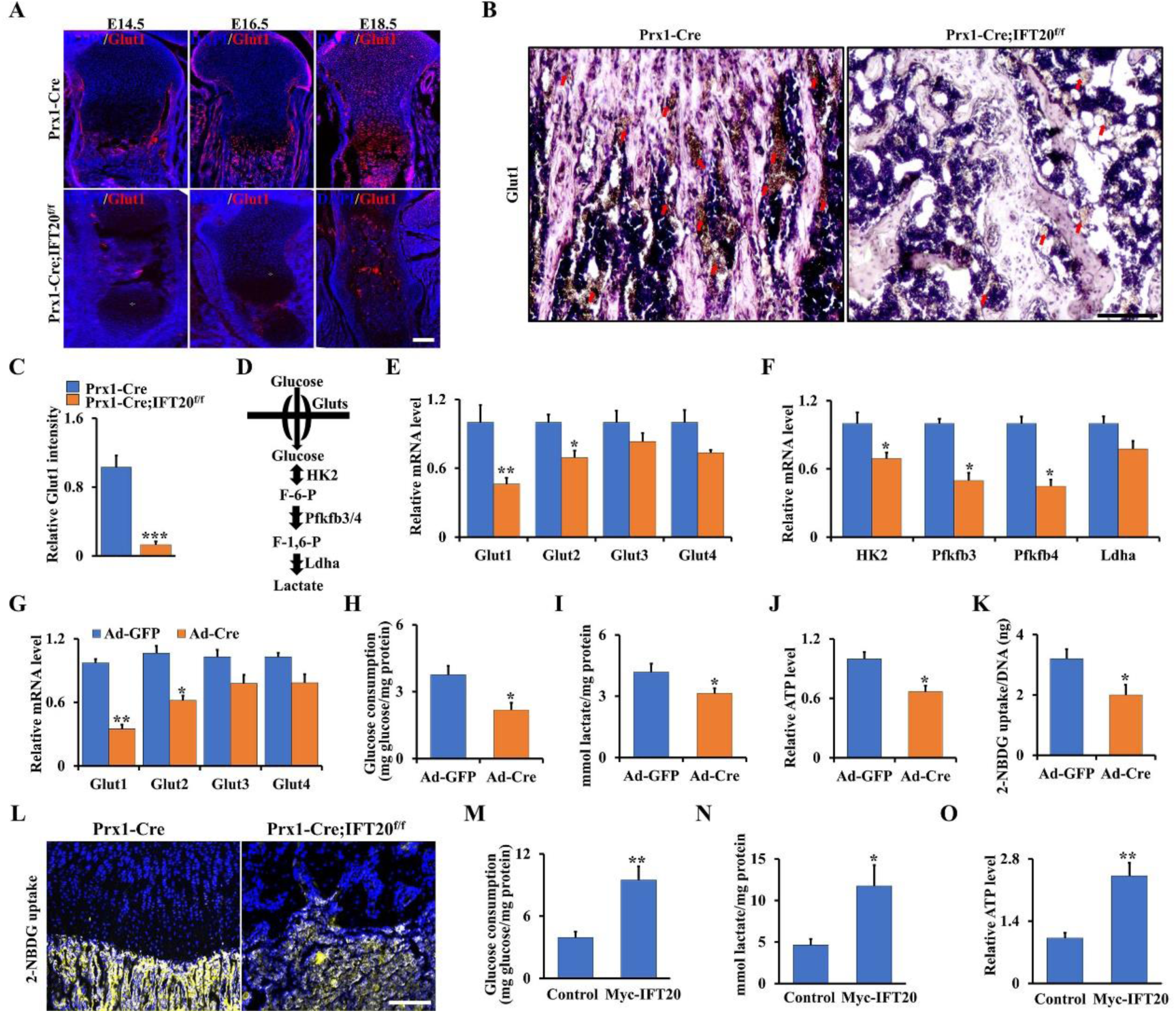
IFT20 promotes glucose uptake and glycolysis in MSCs through Glut1 signaling. (A) Representative image of immunofluorescence staining for Glut1 in tibiae from Prx1-Cre;IFT20^f/f^ mice and age-mated controls as indicated. Scale bars, 75 μm. (B, C) Representative immunohistochemistry image of Glut1 in tibiae from 1-month-old Prx1-Cre;IFT20^f/f^ mice and controls as indicated. Scale bars, 75 μm. Relative intensity of Glut1 was measured as indicated (C). (D) Diagram of the key enzymes in the glucose metabolism such as Gluts, Hk2, Pfkfb3/4 and Ldha. (E, F) qRT-PCR analysis of glucose metabolism-related genes (Glut1/2/3/4, Hk2, Pfkfb3/4, and Ldha) using RNA from MSCs from 1-month-old Prx1-Cre;IFT20^f/f^ mice and controls as indicated. (G) mRNA levels of Glut1-4 in MSCs from IFT20^f/f^ mice after treatment with Ad-GFP or Ad-Cre for 48 hr. (H, I) Glucose consumption and lactate production after treatment of Ad-GFP or Ad-Cre for 48 hr as indicated. (J) ATP production. (K) 2-NBDG uptake. After incubation with 100 μM 2-NBDG of 8 hr, the glucose uptake was determined by a fluorescence microscope at 485/540 nm. (L) Visualization of 2-NBDG uptake in tibiae after injection of 2-NBDG for 45 minutes. (M-O) The glucose consumption (M), lactate production (N), and ATP level (O) were identified after overexpression of IFT20 in MSCs for 48 hr. Error bars were the means ± SEM from three independent experiments. \**P* < 0.05, \*\**P* < 0.01.

### IFT20 promotes Glut1 expression through TGF-β-Smad2/3 signaling

Previous evidence showed that TGF-β signaling is a critical regulator of bone formation and involved in glucose metabolism through regulation of Glut1 expression (Kitagawa et al. 1991; Andrianifahanana et al. 2016; Wu et al. 2016; Xu et al. 2018). Additionally, our previous findings also revealed that loss of IFT80 in chondrocytes inhibited chondrogenesis and fracture healing through inhibiting TGF-β-Smad2/3 signaling (Liu et al. 2020). To get further insight into regulation of IFT20 in glucose metabolism and skeletal development, we isolated total RNA from IFT20-deficient osteoblast progenitor cells and controls, respectively, and performed RNA-seq. KEGG analysis suggested TGF-β signaling pathway had a significant change after loss of IFT20 (Fig. 7A). Moreover, GSEA also showed the signature genes for TGF-β signaling were notably downregulated due to loss of IFT20 (Fig. 7B), as evidenced by pSmad2/3 expression and nuclear location (Fig. 7C, D). To further identify the regulation of Smad2/3 in Glut1, we analyzed the DNA binding motif (CAGAC) of Smad2/3 in the *Glut1* promoter using Vector NTI software. As expected, we found there were four DNA binding sites of Smad2/3 in the *Glut1* promoter (Fig. 7E). Next, we performed the ChIP assay using MSCs isolated from Prx1-Cre;IFT20^f/f^ mice and controls. The specific DNA binding regions of Smad2/3 was amplified by qRT-PCR after immunoprecipitation with Smad2/3 antibody. Our data showed that the transcriptional activity of the *Glut1* promoter significantly decreased after loss of IFT20 (Fig. 7F). Moreover, we also found that Glut1 stability was enhanced after overexpression of IFT20 (Fig. 7G). Taken together, our results suggested that IFT20 governs glucose metabolism through TGF-β-Smad2/3-Glut1 axis.

**Figure 7.**
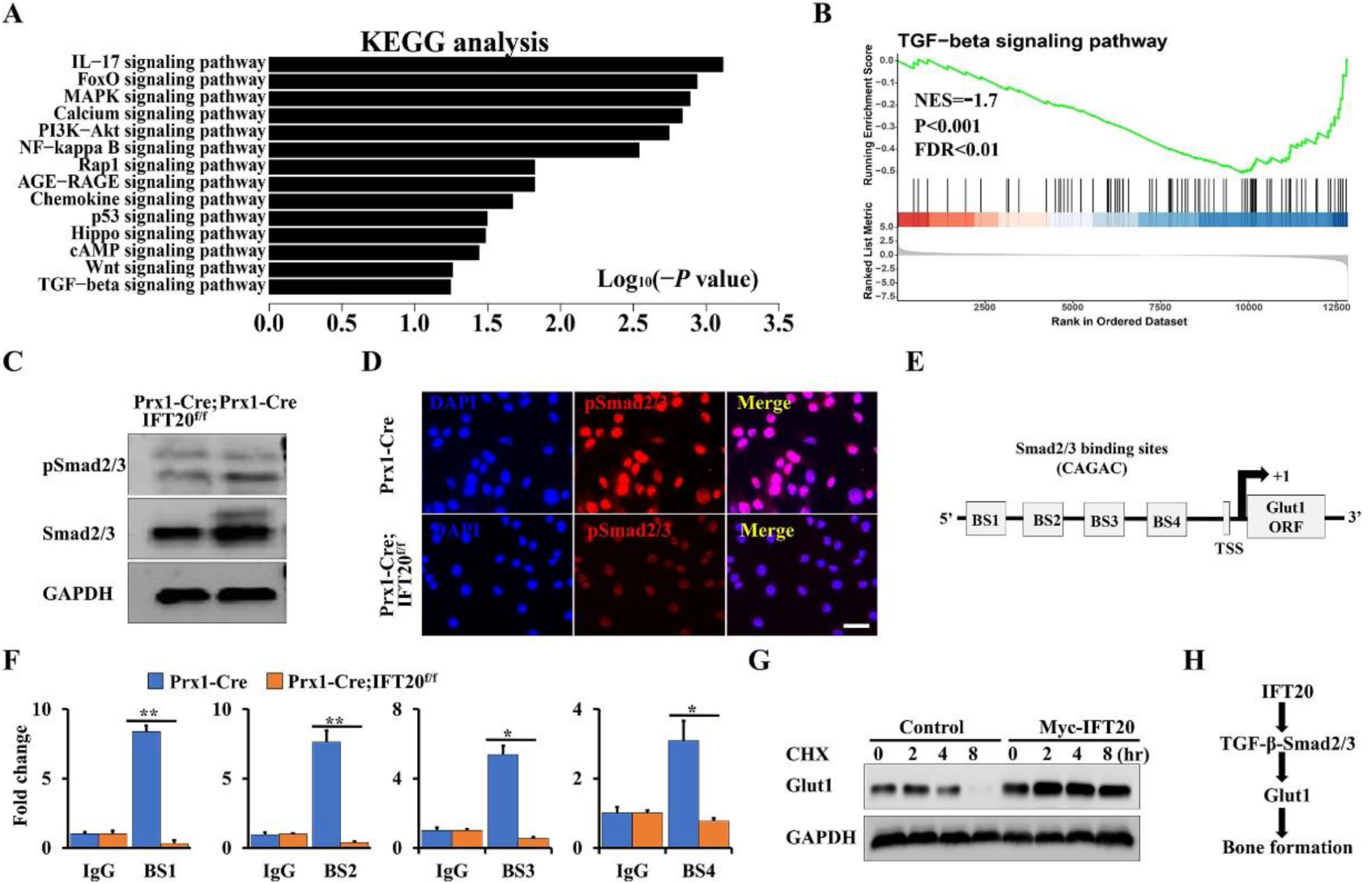
IFT20 promotes Glut1 expression through TGF-β-Smad2/3 signaling. (A) KEGG analysis of significant change of genes after loss of IFT20 in MSCs. (B) GSEA analysis showed a significant decrease of TGF-β signaling after loss of IFT20 in MSCs as indicated. NES, normalized enrichment score. FDR, false discovery rate. (C) western blot analysis as shown. (D) Representative fluorescence image of pSmad2/3 as shown. Scale bars, 10 μm. (E) Schematic diagram of Smad2/3 DNA binding motifs in the *Glut1* promoter. BS, binding site. TSS, transcription start site. (F) ChIP assay. Co-occupation of Smad2/3 in the *Glut1* promoter as indicated. (G) After transfection of Myc-IFT20 or empty vector for 24 hr in MSCs, the MSCs were treated with 50 μg/mL cycloheximide (CHX) for different times as indicated, and then the Glut1 levels were identified by western blot. (H) Proposed mechanism of IFT20 governs mesenchymal stem cell fate through TGF-β-Smad2/3-Glut1 axis. IFT20 in MSCs favors to osteogenesis instead of adipogenesis by maintaining the expression and stability of TGF-β-Smad2/3-mediated Glut1 and enhancing its mediated glucose metabolism. Error bars were the means ± SEM from three independent experiments. \**P* < 0.05, \*\**P* < 0.01.

## Discussion

Aberrant lineage allocation of MSCs contributes to marrow osteoblast-adipocyte imbalance. However, little is known about what the signaling molecules regulate this balance. Here, by genetic deleting IFT20 in a mesenchymal line using Prx1-Cre, we found for the first time that loss of IFT20 in MSCs causes severe limb shorten and bone loss accompanied by significantly increased MAT accumulation in Prx1-Cre;IFT20^f/f^. Mechanistically, we revealed that IFT20 is a positive and key regulator of Glut1 expression and stability as well as glucose metabolism. Loss of IFT20 in MSCs impaired glucose metabolic homeostasis and thereby disrupted the balance of osteoblast and adipocyte differentiation from MSCs. Thus, this study revealed IFT20 as a new and critical regulator of MSC lineage commitment, suggesting that IFT20 may be a promising drug target for treatment of bone diseases such as osteoporosis and other metabolic diseases.

Accumulating evidence indicates that Prx1-Cre is expressed in mesenchymal progenitors throughout skeletal development, including both the embryonic and adult stages, while Lepr-Cre is mainly expressed in adult stem cells (Yu et al. 2018; Yu et al. 2019; Deng et al. 2021). Here, we found that IFT20 deficiency in Prx1^+^ cells caused bone loss and MAT accumulation; however, IFT20 deficiency in Lepr^+^ cells did not result in a statistically significant difference in adipocyte numbers even though bone mass was also decreased compared to that in the age-matched controls, suggesting that IFT20 facilitates MSC lineage commitment at the early stage of MSC differentiation. Our previous study showed that deletion of IFT20 in the osteoblast lineage using OSX-Cre and Collagen-Cre^ERT^ resulted in a reduction in bone mass (Lim et al. 2020). More importantly, we did not find any pronounced phenotype after deletion of IFT20 in osteocytes using DMP1-Cre, reinforcing our finding that IFT20 regulates bone homeostasis at the early stage of skeletal development rather than at the late stage. Similar to our findings, deletion of PTH1R at the early stage of MSCs regulates mesenchymal cell fate, however, when PTH1R was deleted in osteoprogenitor stage, there was no effect on MSC lineage commitment (Fan et al. 2017). Additionally, loss of glutaminase (Gls) in Prx1^+^ cells caused impaired bone mass accompanied by excessive MAT accumulation in the bone marrow; however, no fat change in bone marrow was detected in Lepr-Cre;Gls^f/f^ mice (Yu et al. 2019). Of note, we found that conditional deletion of IFT20 in adipocytes by Adipoq-Cre increased the bone mass. These findings strongly supported the tenet that IFT20 governed early stage MSCs lineage commitment.

Previous studies have demonstrated that glucose metabolism regulates skeletal development by enhancing glucose uptake and glycolysis (Matsumoto et al. 2017; Dirckx et al. 2018; Karner and Long 2018; Wang et al. 2021). Moreover, cilia-related proteins have been proven to be important for maintaining energy balance (Han et al. 2014; Lee et al. 2015; Song et al. 2018). For instance, the deletion of IFT88 in pancreatic β-cells impaired glucose homeostasis and further led to the development of diabetes (Volta et al. 2019; Hughes et al. 2020). Additionally, increased glucose was observed in hyperphagia-induced obesity caused by global deletion of IFT88 or Kif3a in tamoxifen-inducible CAGG-Cre^ERT^ (Davenport et al. 2007; Oh et al. 2015). Interestingly, we found that deletion of IFT20 in MSCs decreased Glut1 expression and increased blood glucose levels, but did not increase insulin sensitivity with high insulin production in either the Prx1-Cre^ERT^;IFT20^f/f^ mice or the Prx1-Cre;IFT20^f/f^ mice, suggesting that IFT20 governs MSCs’ fate by enhancing glucose uptake and metabolism instead of a reduction in serum insulin. It’s well-known that TGF-β signaling is a critical regulator of bone formation and glucose metabolism through regulation of Glut1 expression (Kitagawa et al. 1991; Andrianifahanana et al. 2016; Wu et al. 2016; Xu et al. 2018). Moreover, we previously also found IFT80 promotes bone formation and fracture healing through enhancing TGF-β-Smad2/3 pathway (Liu et al. 2020). In consistent with those findings, our RNA-seq data reveled that TGF-β-Smad2/3 pathway was significantly prohibited due to IFT20 deficiency. Moreover, we found that IFT20 drives glucose uptake and consumption by regulating the expression and stability of TGF-β-Smad2/3-mediated Glut1. These results were further supported by the findings that loss of Vhl in osteoblast progenitor cells enhanced bone mass by improving global glucose metabolism (Dirckx et al. 2018); in contrast, an impaired glucose metabolism led to a significant bone loss (Matsumoto et al. 2017; Lee et al. 2018). Lee *et al*. reported that serum OCN is negatively associated with plasma glucose (Lee et al. 2007; Dirckx et al. 2018). In line with this, we found a marked decrease in serum OCN and increased blood glucose after deletion of IFT20 in MSCs. Correspondingly, Glut1-mediated glucose metabolism has been demonstrated to directly regulate skeletal development in Prx1-Cre;Glut1^f/f^ mice (Lee et al. 2018).

Bone formation is tightly regulated by adipocytes as well as osteoclasts and osteoblasts. Recent studies have highlighted the fact that bone marrow adipocytes can secrete RANKL and support osteoclast differentiation (Kuhn et al. 2012; Fan et al. 2017). Yu *et al*. reported that bone marrow adipogenic lineage precursors inhibit bone formation and promote bone resorption (Yu et al. 2021). Interestingly, our TRAP staining results showed a significant increase in osteoclastogenesis after deletion of IFT20 in both Prx1^+^ and Lepr^+^ cells. Furthermore, when we deleted IFT20 in adipocytes using Adipoq-Cre, Adipoq-Cre;IFT20^f/f^ mice displayed a phenotype of increased bone mass with a significant decrease in osteoclast numbers. Importantly, the expression and content of RANKL in the serum were significantly upregulated in whole bone marrow and bone marrow adipose tissue after deletion of IFT20 in MSCs. Conversely, IFT20 deficiency in adipocytes reversed these phenotypes. Notably, we also found that many adipocytes were present in very close proximity to the bone surface and covered by TRAP^+^ osteoclasts in the Prx1-Cre;IFT20^f/f^ mice and observed a remarkably increased ratio of RANKL/OPG in the serum of the Prx1-Cre;IFT20^f/f^ mice compared to the control mice, suggesting that adipocyte-derived RANKL is the principal source for bone resorption in the absence of IFT20 signaling. These findings were supported by a previous report that RANKL levels were increased after deletion of PTH1R in MSCs in bone marrow and bone marrow adipose tissue, and adipocytes were the most important source of RANKL in the absence of PTH1R signaling (Fan et al. 2017).

In summary, this study uncovers IFT20 as a new regulator governs MSC lineage commitment and balances between osteogenesis and adipogenesis through regulating glucose metabolism. This study provides valuable insights for developing novel therapeutic targets for osteoporosis, obesity, and other ciliopathies.

## Materials and methods

### Animals

Prx1-Cre, Lepr-Cre, DMP1-Cre, and IFT20^f/f^ mice were ordered from The Jackson Laboratory (Bar Harbor, MA, USA). Prx1-Cre^ERT^ mice were a gift from Dr. Dana Graves’s lab at the School of Dental Medicine, University of Pennsylvania. Adipoq-Cre mice were a gift from Dr. Ling Qin’s lab at Perelman School of Medicine, University of Pennsylvania. For the Prx1-Cre^ERT^;IFT20^f/f^ and Prx1-Cre^ERT^ mice, tamoxifen was administered at D9 and D13.

### Antibodies, reagents and plasmids

Antibodies against Glut1 and TRAP staining kits were ordered from Sigma. Antibodies against MMP13, BrdU and ColX were obtained from Santa Cruz Biotechnology. The secondary fluorescent antibodies and H&E staining kit were from Abcam. The fluorescent glucose analog 2-(N-(7-nitrobenz-2-oxa-1,3-diazol-4-yl) amino)-2-deoxyglucose (2-NBDG) was purchased from Cayman Chemical Company. BrdU labeling and calcein labeling reagents were purchased from Fisher Scientific™. The plasmids pcDNA3.1-Myc and pcDNA3.1-Myc-IFT20 were obtained from Addgene. The transfection reagents (FuGENE^®^ HD) were from Promega Corporation.

### Cell culture

Primary MSCs were isolated from the femurs of Prx1-Cre;IFT20^f/f^ mice and other corresponding controls. Briefly, the fresh femurs were thoroughly cleaned after removing all soft tissues, especially muscles, and then, the bone marrow was thoroughly flushed out with α-minimum essential medium (α-MEM; Gibco, USA), collected and cultured in α-MEM supplemented with 10% fetal bovine serum (FBS) (Gibco, USA) and 1× Pen-Strep solution (Fisher Scientific™, USA) at 37 °C with 5% humidified CO_2_. The medium was replaced every other day.

### Colony formation unit assay

For the CFU assay, briefly, 5×10^3^ primary MSCs isolated from the femurs of Prx1-Cre;IFT20^f/f^ mice and controls were seeded in 6-well plates and cultured in regular α-MEM medium at 37 °C with 5% humidified CO_2_. After incubation for 5 days, the 6-well plate was stained with 0.5% crystal violet, and then, the colony numbers were counted in three different plates.

### ALP staining, ALP activity, and osteoblast differentiation

For ALP staining, briefly, primary MSCs from Prx1-Cre;IFT20^f/f^ mice and controls were induced in α-MEM containing 10% FBS, 1× Pen-Strep solution, 100 nM dexamethasone (Sigma, USA), 50 μg/mL L-ascorbic acid (Sigma, USA), and 5 mM β-glycerophosphate (Sigma, USA). The osteogenic medium was replaced every 3 days. After 5 days of osteogenic induction, the cells were fixed with PFA for 30 seconds at room temperature, and then, ALP staining was performed by a BCIP/NBT ALP Staining Kit (Millipore, USA) according to the manufacturer’s instructions.

After 7 days of osteogenic induction, ALP activity was measured by a microplate reader at OD_405_ nm. Briefly, cells were harvested with harvest buffer (2 mM PMSF and 0.2% NP-40 in 10 mM Tris-Cl (pH 7.4)) after washing 2 times with ice-cold PBS, and then, the supernatants were collected and incubated with assay buffer containing 1 mM MgCl_2,_ 100 mM glycine (pH 10.5) and 50 mM p-nitrophenyl phosphate solution for 15 min at 37 °C. Subsequently, the reaction was stopped by 0.1 N NaOH solution, the samples were assayed at OD_405_ nm, and the production of p-nitrophenol (nmol) in total protein (per min per mg) was determined.

Osteogenic differentiation was induced for 2 weeks by osteogenic medium as we described above, and then, staining with Alizarin Red S staining solution (pH 4.4) was performed. After the images were scanned, the stains were thoroughly destained by 10% cetylpyridinium chloride in 10 mM sodium phosphate buffer (pH 7.0), measured and quantified by a microplate reader at OD_562_ nm.

### Oil red O staining

Adipogenic differentiation was induced for 2 weeks by adipogenic medium (α-MEM supplemented with 10% FBS, 1×Pen-Strep solution, 10 μg/mL insulin, 100 μM rosiglitazone, 500 μM 3-isobutyl-1-methylxanthine (IBMX), and 1 μM dexamethasone) and then stained with Oil red O solution. After these images were scanned, the stained plates were thoroughly washed with isopropanol, measured and quantified by a microplate reader at OD_405_ nm.

### Metabolite measurements

MSCs were isolated from femurs of Prx1-Cre;IFT20^f/f^ mice and controls or IFT20 floxed mice as outlined in Cell Culture. Each cell pool was split into a pair of a control group and an experimental group and then transduced with adenovirus Ad-Cre or control Ad-GFP. Next, glucose consumption and lactate production were measured by the Glucose (HK) Assay Kit (Sigma, USA) and the L-lactate Assay Kit (Eton Biosciences, USA), respectively, according to the manufacturer’s instructions. For the glucose uptake assay, after incubation with 100 μM 2-NBDG for 8 hr, glucose uptake was determined by a fluorescence microscope at 485/540 nm. ATP production was quantified using the CellTiter-Glo^®^ Luminescent Cell Viability Assay kit (Promega, USA). Serum levels of OCN, OPG, RANKL and insulin were measured by mouse Osteocalcin ELISA Kit (BioVision), OPG ELISA Kit (Boster Biological Technology), TNFSF11/RANKL PicoKine ELISA Kit (Boster Biological Technology) and Ultra-Sensitive Insulin ELISA Kit (Crystal Chem), respectively, according to the manufacturer’s instructions. PAS staining was carried out by a periodic acid-Schiff (PAS) kit (Sigma, USA) according to the manufacturer’s instructions.

Bone marrow adipose tissue from Prx1-Cre;IFT20^f/f^ mice, Adipoq-Cre;IFT20^f/f^ mice and corresponding controls was isolated as described previously (Fan et al. 2017). Briefly, the fresh femurs were thoroughly cleaned after removing all soft tissues, and then, the bone marrow was quickly flushed out with α-MEM medium in 1.5 mL EP tubes. The cells were collected by centrifugation, and red blood cell lysis buffer was added for incubation. Then, floating adipocytes were collected from the top layer by centrifugation for 5 min at 3,000 rpm.

### Blood glucose test and 2-NBDG tracing

For the glucose tolerance test, the mice were fasted overnight and then injected intraperitoneally with sterile glucose (2 g/kg body weight) as previously reported (Matsumoto et al. 2017; Dirckx et al. 2018). The blood glucose level was monitored by a blood glucose meter (OneTouch® Ultra^®^2). Then, 2-NBDG tracing was conducted in 1-month-old Prx1-Cre;IFT20^f/f^ mice and controls. Briefly, anesthetized mice were injected with 25 mg/kg 2-BDNG via the tail vein. After injection for 45 minutes, the mice were sacrificed and fixed in 4% paraformaldehyde (PFA) overnight at 4 °C. Then, the tibiae were decalcified with 10% EDTA in PBS (pH 7.4) for 2 weeks, embedded in OCT, sectioned, and analyzed by a fluorescence microscope (Dirckx et al. 2018).

### Immunofluorescence and immunohistochemistry

For immunofluorescence staining, tibial sections collected at P0 were prepared in advance. Briefly, the dehydrated tibiae were dehydrated through serial incubations of ethanol and xylene and then embedded in paraffin. Next, the sections were blocked with 1% BSA for 1 hr at room temperature and probed with primary antibodies against MMP13 (1:200 dilution), ColX (1:200 dilution), BrdU (1:200 dilution), and Glut1 (1:100 dilution) overnight at 4 °C. After 3 washes with PBST, the cells were incubated with the corresponding secondary fluorescent antibody (1:1000 dilution) for 1 hr in the dark. Then, counterstaining of nuclei was performed with DAPI, followed by 3 washes with PBST and visualization under a fluorescence microscope as we previously reported (Li et al. 2021a; Li et al. 2021b). Immunohistochemistry was carried out as we previously reported (Li et al. 2021a).

### qRT-PCR, ChIP-qPCR and western blot

Briefly, 1 μg total RNA extracted from the cortical bones of 1-month-old Prx1-Cre, Prx1-Cre;IFT20^f/f^, and IFT20 floxed mice using TRIzol reagent (TaKaRa, Japan) was reverse-transcribed into cDNA by PCR using PrimeScript™ RT Kit (TaKaRa, Japan). Then, qRT-PCR was performed using a CFX96 Real-Time PCR System and SYBR Green mixture (Bio-Rad, USA). GAPDH served as an internal control and was determined by the 2^−ΔΔCt^ method. ChIP-qPCR and western blotting were carried out as we previously reported (Li et al. 2021a; Li et al. 2021b). The primers of qRT-PCR and ChIP-qPCR used in this study were listed in Supplementary Table S1.

### Whole-mount skeletal staining

Prx1-Cre;IFT20^f/f^ mice and age-matched controls at the indicated timepoints were euthanized and fixed in 100% ethanol at room temperature, and then, whole-mount skeletal staining was carried out as we previously reported (Li et al. 2021b).

### Calcein labeling and histology

Calcein (20 mg/kg) was injected on Day 2 and Day 5 before 1-month-old Prx1-Cre;IFT20^f/f^, Lepr-Cre;IFT20^f/f^ mice and controls were sacrificed. After sacrifice, the tibiae were collected, fixed in 4% PFA overnight at 4 °C, infiltrated in 10% potassium hydroxide (KOH) for 3 days, dehydrated by ethanol and xylene, and then embedded in paraffin. The paraffin sections were prepared at six-micrometer thickness and observed under a fluorescence microscope. The mineral apposition rate (MAR) and bone formation rate per bone surface (BFR) were analyzed by the Leica microanalysis system as we previously reported (Ng et al. 2019; Li et al. 2021b).

For histology, briefly, femurs were harvested, fixed in 4% PFA overnight at 4 °C, decalcified with 14% EDTA in PBS (pH 7.4) for 1 month, and then embedded in paraffin. Six-micrometer sections of these femurs were prepared, and then, H&E and TRAP staining was conducted with an H&E staining kit (Abcam, USA) and TRAP staining kit (Sigma, USA), respectively, as we previously reported (Yuan et al. 2016; Ng et al. 2019; Li et al. 2021b).

### Radiographic analysis

The bone morphology and microarchitecture of femurs from 1-month-old Prx1-Cre;IFT20^f/f^, Lepr-Cre;IFT20^f/f^, Adipoq-Cre;IFT20^f/f^, and DMP1-Cre;IFT20^f/f^ mice and controls were analyzed by a high-resolution micro-CT system (images acquired at 55 kV energy, 145 mA, and 300 ms integration time) as we described previously (Ng et al. 2019; Lim et al. 2020; Li et al. 2021b).

### RNA-seq and bioinformatic analysis

RNA-sequencing was performed as previously described (Tang et al. 2021). Briefly, total RNA was isolated using TRIzol reagent from IFT20-deficient osteoblast progenitor cells and controls, respectively. And then the library and sequencing were carried out by the RNA-Seq Core at University of Pennsylvania. Read counts were subjected to paired differential expression analysis using the R package DESeq2.

### Statistical analysis

In this study, other analyses of experimental data were conducted and analyzed by SPSS 21 software, and data are reported as the mean ± SEM by Student’s t test. The statistical significance of group differences was determined by 2-way ANOVA. *P* values < 0.05 were considered significant.

## Acknowledgments

We are thankful to the animal facility at University of Pennsylvania. This study was supported by the National Institute of Arthritis and Musculoskeletal and Skin Diseases (NIAMS, AR061052), the National Institute of Aging (NIA, AG048388) and the National Institute of Dental and Craniofacial Research (NIDCR, DE023105) awarded to Shuying Yang.

## Author Contributions

Shuying Yang and Yang Li conceived this study, generated hypotheses, and designed experiments. Yang Li, Shuting Yang and Yang Liu performed experiments and analyzed data. Shuying Yang, Yang Li and Ling Qin wrote, reviewed and edited the paper. Shuying Yang supervised the project.

## Data and materials availability

The data that support the findings of this study are available in the paper and supplementary materials.

