## supplemental information for "IFT20 governs mesenchymal stem cell fate through positively regulating TGF-β-Smad2/3-Glut1 signaling mediated glucose metabolism"

**Supplemental Figure Legends**

Figure S1. IFT20 deficiency in Prx1-expressing cells causes a severe limb shortening in mice. (A) IFT20 expression. qRT-PCR analysis using RNA from MSCs from Prx1-Cre;IFT20$^{f/f}$ mice and controls. (B) Representative whole-mount skeletal-stained image of Prx1-Cre;IFT20$^{f/f}$ mice and age-mated controls at different timepoints of embryo (E16.5 and E18.5) as indicated. (C) Quantitative analysis of the length of femurs and tibiae from Prx1-Cre;IFT20$^{f/f}$ mice and age-mated controls as indicated. (D, E) Representative images of Prx1-Cre;IFT20$^{f/f}$ mice and controls at age of 1 month. (F) Quantitative analysis of the length of femurs and tibiae from 1-month-old Prx1-Cre;IFT20$^{f/f}$ mice and controls. Error bars were the means ± SEM from three independent experiments. *$P < 0.05$, **$P < 0.01$, ***$P < 0.001$.

Figure S2. IFT20 deficiency in Prx1-expressing cells impaired bone formation. (A) Representative micro-CT image of femurs of Prx1-Cre;IFT20$^{f/f}$ mice and age-matched controls at P7. Scale bars, 1 mm. (B) Representative micro-CT image of femurs of Prx1-Cre;IFT20$^{f/f}$ mice and age-matched controls at 1 month. Scale bars, 1 mm.

Figure S3. IFT20 deficiency in Lerp-expressing cells causes bone-fat imbalance. (A) Representative micro-CT image of femurs of Lepr-Cre;IFT20$^{f/f}$ mice and controls at 1 month. Scale bars, 1 mm. (B-E) Histomorphometric analysis of bone parameters in the femurs of 1-month-old Lepr-Cre;IFT20$^{f/f}$ mice and controls. Bone volume fraction (BV/TV); trabecular thickness (Tb.Th); trabecular number (Tb.N); trabecular spacing (Tb.Sp). N=5 mice/group. (F) Quantitative measurements of BMD of femurs from Lepr-Cre;IFT20$^{f/f}$ mice and controls at 1 month. (G) The serum level of OCN from Lepr-Cre;IFT20$^{f/f}$ mice and controls at 1 month. (H-J) Calcein double labeling in tibia of 1-month-old Lepr-Cre;IFT20$^{f/f}$ mice and controls. Scale bar, 50 μm. (K) Representative H&E-stained image of femur sections from Lepr-Cre;IFT20$^{f/f}$ mice and controls at 1 month. Scale bars, 200 μm. (L) Representative TRAP-stained image of femur sections from 1-month-old Lepr-Cre;IFT20$^{f/f}$ mice and controls. The corresponding quantitative analysis was performed at lower panel. (M) $OsO_4$ micro-CT staining of decalcified tibiae by micro-CT analysis as indicated. Error bars were the means ± SEM from three independent experiments. *$P < 0.05$, **$P < 0.01$.

Figure S4. The effect of IFT20 on mature osteoblasts is dispensable. (A) Representative micro-CT image of femurs of DMP1-Cre;IFT20$^{f/f}$ mice and controls at 1 month. Scale bars, 1 mm. (B) Histomorphometric analysis of bone parameters in the femurs of 1-month-old DMP1-Cre;IFT20$^{f/f}$ mice and controls.

Figure S5. IFT20 expression. (A) IFT20 expression was identified by qRT-PCR after transfection for 48 hr with Ad-Cre or Ad-GFP in the MSCs from IFT20$^{f/f}$ mice. (B) MSCs were transfected with Myc-IFT20 plasmid or empty vector, respectively. After transfection of 48 hr, the IFT20 expression was identified by qRT-PCR. Error bars were the means ± SEM from three independent experiments. ***$P < 0.001$.

**Figure. S1**

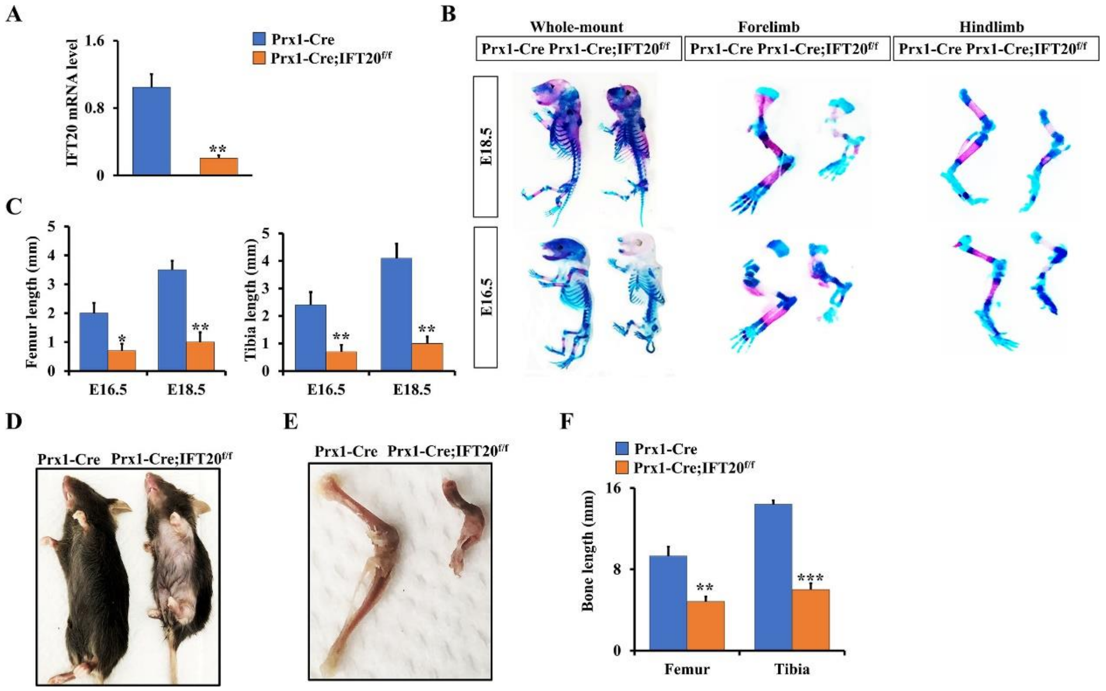

A
IFT20 mRNA level
1.6
0.8
0
**
Prx1-Cre
Prx1-Cre;IFT20f/f
B
Whole-mount
Forelimb
Hindlimb
Prx1-Cre Prx1-Cre;IFT20f/f
Prx1-Cre Prx1-Cre;IFT20f/f
Prx1-Cre Prx1-Cre;IFT20f/f
E18.5
E16.5
C
Femur length (mm)
Tibia length (mm)
5
4
3
2
1
0
*
**
E16.5
E18.5
D
Prx1-Cre Prx1-Cre;IFT20f/f
E
Prx1-Cre Prx1-Cre;IFT20f/f
F
Prx1-Cre
Prx1-Cre;IFT20f/f
Bone length (mm)
16
8
0
**
***
Femur
Tibia

**Figure. S2**

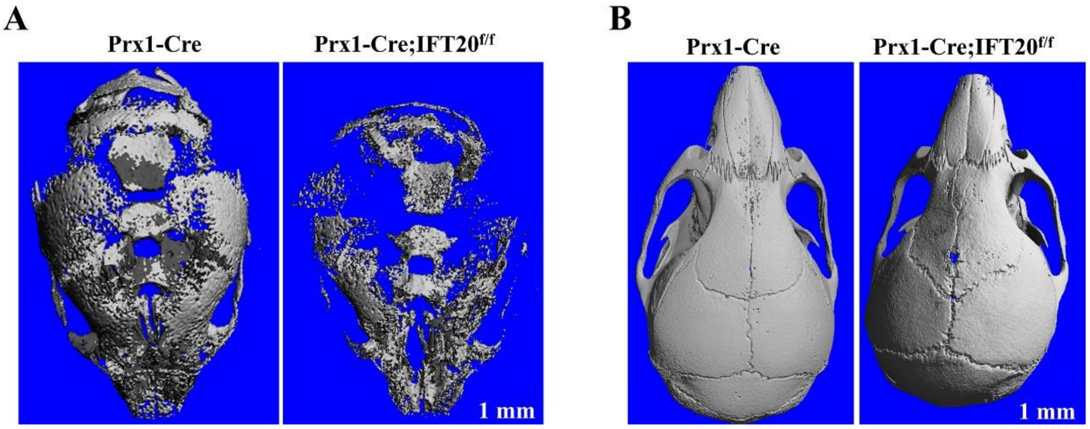
A
Prx1-Cre
Prx1-Cre;IFT20f/f
1 mm
B
Prx1-Cre
Prx1-Cre;IFT20f/f
1 mm

**Figure. S3**

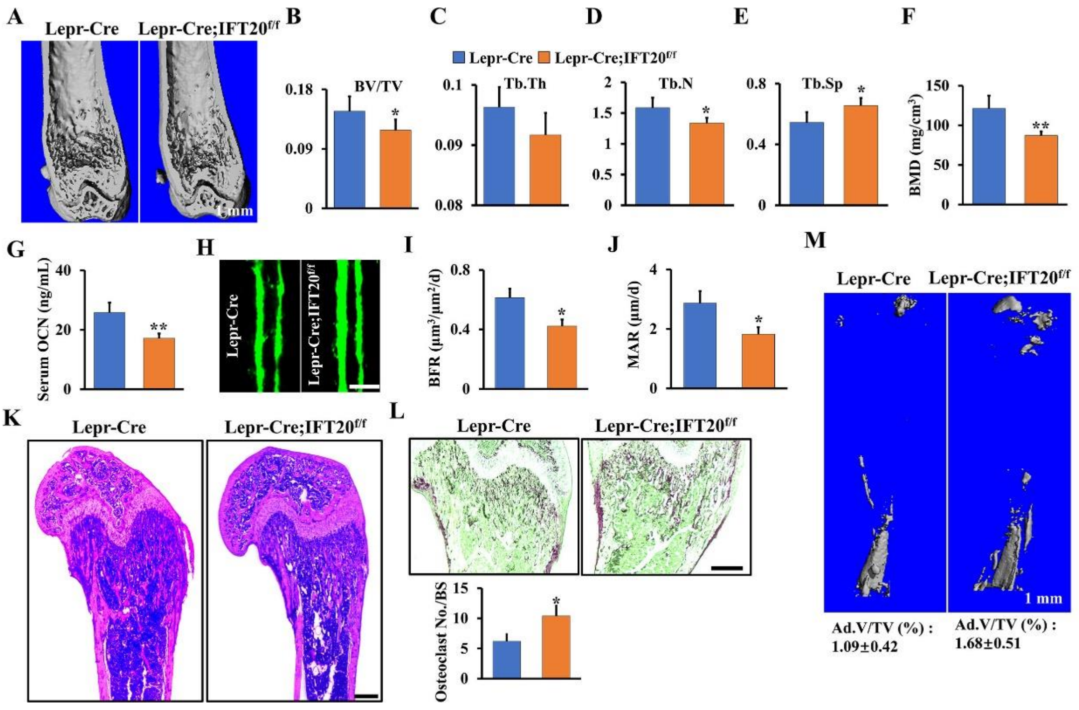
A
Lepr-Cre
Lepr-Cre;IFT20f/f
1 mm
B
C
D
E
F
Lepr-Cre
Lepr-Cre;IFT20f/f
BV/TV
Tb.Th
Tb.N
Tb.Sp
BMD (mg/cm3)
G
Serum OCN (ng/mL)
H
Lepr-Cre
Lepr-Cre;IFT20f/f
I
BFR (μm3/μm2/d)
J
MAR (μm/d)
M
Lepr-Cre
Lepr-Cre;IFT20f/f
1 mm
Ad.V/TV (%) : 1.09±0.42
Ad.V/TV (%) : 1.68±0.51
K
Lepr-Cre
Lepr-Cre;IFT20f/f
L
Lepr-Cre
Lepr-Cre;IFT20f/f
Osteoclast No./BS

133 **Figure. S4**

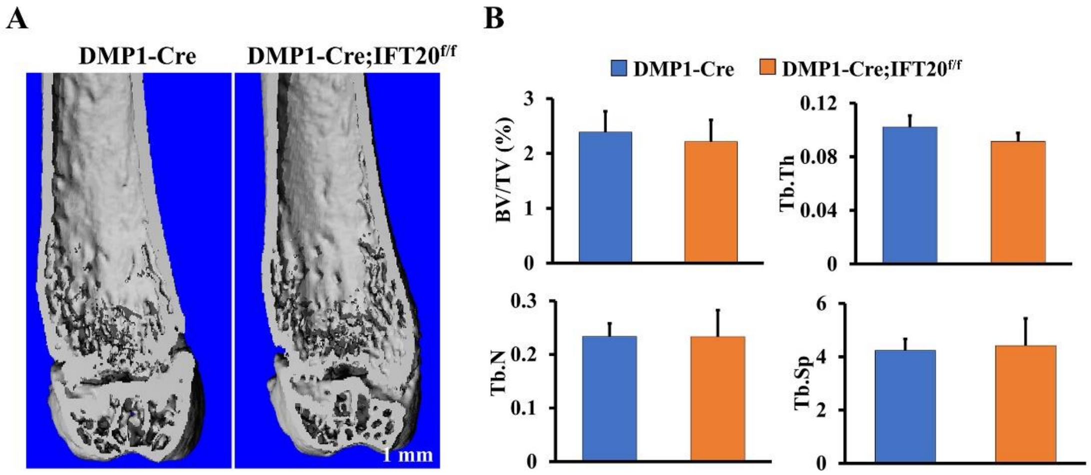

**Figure. S5**

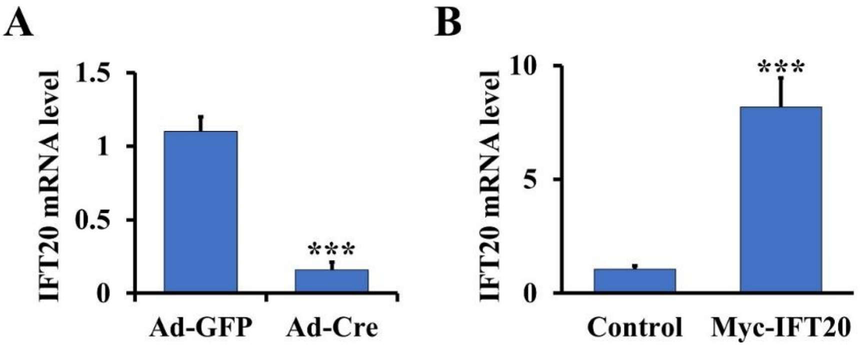
A
IFT20 mRNA level
1.5
1
0.5
0
***
Ad-GFP
Ad-Cre
B
IFT20 mRNA level
10
5
0
***
Control
Myc-IFT20

### Supplementary information, Table S1

| Gene | sequence （5'-3'） | Gene | sequence （5'-3'） |
|---|---|---|---|
| GAPDH-F | CCTGGTCACCAGGGCTGCCATTT | Runx2 | GCCGGGAATGATGAGAACTA |
| GAPDH-R | CGTTGAATTTGCCGTGAGTGGAG | Runx2 | GGACCGTCCACTGTCACTTT |
| Glut2-F | ATCCCTTGGTTCATGGTTGCTG | ALP-F | AAGGCTTCTTCTTGCTGGTG |
| Glut2-R | TCCGCAATGTACTGGAAGCAG | ALP-R | GCCTTACCCTCATGATGTCC |
| Glut3-F | TGGTAGCTCAGATCTTTGGTTTGG | OSX-F | GGAGGCACAAAGAAGCCATACGC |
| Glut3-R | GATCTCTGTAGCTTGGTCTTCCTC | OSX-R | TGCAGGAGAGAGGAGTCCATTG |
| Glut4-F | CCAGCCACGTTGCATTGTA | OCN-F | CTTGGTGCACACCTAGCAGA |
| Glut4-R | ACACTGGTCCTAGCTGTATTCT | OCN-R | ACCTTATTGCCCTCCTGCTT |
| IFT20-F | GCGAGGCAGGGCTGCATTTTGAT | Glut1-F | GGGCATGTGCTTCCAGTATGT |
| IFT20-R | CCTTGCACTCCTCCTTGAGCTCC | Glut1-R | ACGAGGAGCACCGTGAAGAT |
| HK2-F | TGATCGCCTGCTTATTCACGG | Pfkfb3-F | CTCCCAGCCCGGGGTAAGACTTACA |
| HK2-R | AACCGCCTAGAAATCTCCAGA | Pfkfb3-R | GCTTCACAGCCTCACGCCGATA |
| Pfkfb4-F | CCGACACTCATTGTCATGGTGG | Ldha-F | TGTCTCCAGCAAAGACTACTGT |
| Pfkfb4-R | CACGCCAATCCAGTTGAGGTAC | Ldha-R | GACTGTACTTGACAATGTTGGGA |
| PPARr-F | ACAGAGATGCCATTCTGGCCCACCAAC | Fabp4-F | ATGTGTGATGCCTTTGTGGGAACC |
| PPARr-R | GCTGGAGAAATCAACTGTGGTAAAGGGC | Fabp4-R | CCATGCCTGCCACTTTCCTTGTG |
| C/EBPa-F | CCGGTGCGGGCAAAGCCAAGAAG | Adiponectin-F | ATGCTACTGTTGCAAGCTCTCCTG |
| C/EBPa-R | TCTTGCGCACCGCGATGTTGTTG | Adiponectin-R | AGGGACCAAAGCAGGAGCTAGCT |
| ChIP1-F | CCTGAAGCTAGCAACAGACT | ChIP3-F | CTTGGGCACAGGAACACGGA |
| ChIP1-R | TCACCAATCAGCCATCTTTT | ChIP3-R | GGTGTTTACAACCGCGTGTG |
| ChIP2-F | GGCAAAGTGGTGATCAGGAG | ChIP4-F | CTGGGACTGCAGGTTCTAGC |
| ChIP2-R | CTAATTGAGCATGGACCCCT | ChIP4-R | TCTGAGAGGCGTGGTTCTGT |
